# Modification of T cells function after restoration of spontaneous circulation in a rat model of cardiac arrest

**DOI:** 10.1101/560789

**Authors:** Chunlin Xing, Yang Chen, Xuemei Zhu, Guoping Lu, Weiming Chen

## Abstract

Cardiac arrest (CA) is a prominent cause of mortality worldwide. A large number of patients after post-cardiac arrest is often associated with a phase of impaired immunity. Through an asphyxial cardiac arrest rat model, we investigate the peripheral blood T cells subsets and the expressions of surface molecules after restoration of spontaneous circulation (ROSC). Sprague-Dawley rats (weight, 300-400 g) were randomly divided into cardiac arrest (CA) group and sham-operated group. CA group rats were induced by 6 minutes of asphyxia. After successful ROSC, 24 surviving rats in two groups were randomly assigned to be sacrificed (n = 8 per subgroup) at 3, 24 and 72 h. The proportion of T cells and CD4+, CD8+ subsets as well as the expression of surface molecules (CTLA-4, PD-1, CD28) on T cells were identified by flow cytometry. The protein concentrations of cytokines (TNF-α, IL-6, IL-10, IL-4, IFN-γ, IL-17A) in serum were measured by ELISA. Compared with sham-operated control group, CD3+ lymphocytes in CA group were significantly decreased at 24 and 72 h post-ROSC. The expression levels of CD28, PD-1, and CTLA-4 on T cells were markedly increased in CA groups at 24 h post-ROSC. Additionally, the concentrations of IFN-γ were significantly declined, while IL-4 was markedly elevated in the CA group at 24 and 72 h post-ROSC. T cells function is moderately changed after CA, which is associated with decreased percentage of T cells, the upregulation of co-inhibitory molecules, and the shift from T helper (Th) 1 to Th2.

## Introduction

Out-of-hospital cardiac arrest (OHCA) is a leading cause of mortality worldwide [1]. Despite many efforts to improve outcomes from sudden cardiac arrest (CA) over past three decades, the survival rate still remains low [2, 3]. During cardiopulmonary resuscitation (CPR) after CA, the body has experienced complete and systemic ischemia, which usually caused the post-cardiac arrest syndrome, a condition characterized by a brain injury, myocardial dysfunction, systemic ischemia/reperfusion response, and persistent precipitating disease [4].

Meanwhile, a large number of cytokines and inflammatory mediators are released in response to the intense pathological stimuli, leading to systemic inflammatory response syndrome (SIRS) [4]. In addition, post-cardiac arrest is often associated with a phase of impaired immunity, during which the patients are more susceptible to secondary infections due to immunosuppression [5, 6]. Furthermore, additional altered immune findings of post-ROSC have been demonstrated, for example, the leukocytes, including T lymphocytes, from post-cardiac arrest patients have been shown to be capable of producing different inflammatory cytokines [7]. Immunosuppression owing to the reduced expression of monocytic human leukocyte antigen-DR (HLA-DR) was also observed in OHCA patients [8]. Besides, it is illustrated that the expression levels of forkhead/winged helix transcription factor (Foxp3) and interleukin 10 (IL-10) in splenic Treg cells were significantly increased in a porcine CA model [9]. Besides, a shift from the Th (T helper) 2 to Th1 is perceived in the myocardium in a porcine model of CA [10].

As the largest group of lymphocytes in immune system, the T cells in septic patients have been repeatedly described in association with nosocomial infections and an increased risk of death [11, 12]. Hence, there are some studies aimed at improving T cells function to enhance the immunity in sepsis patients [13, 14]. However, the function of T cells, especially the expression of T cells surfaces co-stimulatory molecules after CA still remain unclear. Therefore, in our study, a Sprague-Dawley rat model of ROSC from CA was established by asphyxia, and the numbers, differentiation, subsets and the expression of surface molecules of peripheral blood T cells was detected.

## Materials and methods

### Animals

12 to 16-week-old male and female Sprague-Dawley rats (300-400g; Jie Si Jie Laboratory Animal CO., LTD, Shanghai, China) were used in the experiments. Before the experiment, rats were kept in a 22 to 26°C and 12-h light-dark cycle for one week to adapt to the new environment.

### Animal Protocol

The animal protocols as described below were approved by the Institutional Animal Care and Use Committee of Fudan University.

### Animal preparation

The animals were fasted for 12 hours except free access to water before the experiment. Anesthesia was induced by intraperitoneal injection of 10% Chloral hydrate (3.5 ml/kg) and endotracheal intubation used a Y-shaped catheter (Alcott Biotech Company, Shanghai, China). Following skin sterilization, a cut down was performed in bilateral inguinal and intravascular catheters (24 gauge; B. Braun Medical Industries Sdn. Bhd, Penang, Malaysia) were inserted in the right femoral artery for blood pressure monitoring (MP40; PHILIPS, Holland) and the left femoral vein for drug administration. The heart rate was continuously monitored by standard lead Π of the surface electrocardiograph during the experiments. The rats were mechanically ventilated (tidal volume, 4ml; respiratory rate, 80 min^-1^; fraction of inspired oxygen, 21%) with a rodent ventilator (ALC-V8D; Alcott Biotech Company, Shanghai, China).

### Cardiac arrest and cardiopulmonary resuscitation

Asphyxia CA and cardiopulmonary resuscitation were performed as described in previous literature [15]. Briefly, 10 min after the animal preparation, the trachea catheter was blocked at end-expiratory phase and the asphyxia time lasted for 6 min. During trachea blocking, mean arterial pressure (MAP), heart rate and respiratory rate were carefully observed. The definition of CA was the onset of MAP declines to 20 mmHg. After 6 min of asphyxia, cardiopulmonary resuscitation was immediately performed, which is consisted of resuming mechanical ventilation (tidal volume, 4 mL; fraction of inspired oxygen, 100%; inspiratory/expiratory ratio, 1/1.5; respiratory rate 80 min^-1^; ALC-V8D; Alcott Biotech Company ventilator, Shanghai, China), continuous external chest compressions at a rate of 180 to 200 compressions min^-1^ by one person. The intravenous epinephrine (0.02 mg/kg) and bicarbonate (1.0 mEq/kg) was administered as needed until the development of ROSC. ROSC was defined by MAP above 60 mmHg and spontaneous heart rate was observed, in which both indicators sustained for 10 min. When rats were observed to remain in pulseless electrical activity or asystole after CPR performed for 10 min, they were considered as resuscitation failures.

Following ROSC, rats were nursed and maintained to mechanical ventilation for 3 h. The mechanical ventilator would be disconnected if adequate spontaneous respiration recovered, which was defined by the respiratory rate 70-100 min^-1^ on room air and no decrease of MAP for 10 min after disconnecting mechanical ventilator. Subsequently, intravascular catheters were removed and inguinal wounds sutured. Then endotracheal catheter was removed and rats were returned to cages for feeding for 24 h or 72 h. Fluid (5% dextrose in saline, 1:2, 50 ml/kg) was administered intraperitoneal every 12 h during the observation periods for hydration and nutritional supporting. Aside from asphyxia, CPR and mechanical ventilation, rats in sham-operated group underwent all same procedures as those in the CA group.

### Blood sample

The CA group and sham-operated group were respectively allocated to 3 subgroups (3 h subgroup (n = 8), 24 h subgroup (n = 8) and 72 h subgroup (n = 8)), and cardiac blood samples were obtained from all rats after anesthesia and then euthanized with 10% chloral hydrate overdose. One milliliter whole blood anticoagulated by heparin was obtained for flow cytometry, which was performed immediately after withdrawal. While another 3 ml blood samples were collected on sodium citrate and centrifuged at 2000g for 5 min at 4°C, after which the samples were stored at -80°C for further analysis.

## Measurements

### Surface immunofluorescence of T cells

According to the manufacturer’s recommendations, 100 μl whole blood was stained with CD3-Per-eFluor 710 (eBioscience, CA 92121, USA), CD4-V450 (BD Biosciences, MD, USA), CD8-PE-Cy7 (eBioscience), PD-1-APC (eBioscience), CTLA-4-PE (eBioscience), CD28-FITC (eBioscience) at room temperature for 25 min in a dark chamber. Samples were then lysed by the FACS Lysing solution (BD Biosciences) for 10 min at 37°C. Following washing with 2 ml fetal bovine serum (FBS; BD Biosciences), cells were resuspended in 250 μl phosphate buffered saline (PBS). Cells were analyzed by a BD FACSCanto II Cytometer (BD Biosciences) and lymphocyte gate was based upon the forward scatter (FSC) and side scatter (SSC) properties. T cells were identified as CD3+. Analysis was done by FlowJo analysis software (OR, USA).

### Cytokines

The cytokines such as tumor necrosis factor alpha (TNF)-α, IL-6, IL-10, IL-4, IL-17 and interferon gamma (IFN)-γ were measured in plasma by Enzyme-linked Immunosorbent Assay (ELISA) with commercially available kits (Multi sciences (lianke) biotech, CO., LTD. China).

### Statistical Analysis

The experimental data were analyzed by SPSS 17.0 (SPSS Inc., Chicago, IL, USA). The results are reported as mean ± standard deviation (SD) or median (interquartile range), on the basis of their distribution by Shapiro-Wilk test. Student *t* test or Mann-Whitney test was performed to detect differences between the sham and CA variables. A two-tailed value of P < 0.05 was considered statistically significant.

## Results

### Baseline characteristics of animals

Twenty-four rats in the CA group survived and averagely divided into 3 subgroups (time = 3 h (n = 8), time = 24 h (n = 8), time = 72 h (n = 8)), so did the sham-operated group. Baseline and post-ROSC characteristics of rats were summarized in **Table 1**. No significant difference was observed between two groups in sex, body weight, heart rate, MAP and respiratory rate. In CA groups, the induction time was 226 ± 28 s, CA duration was 136 ± 26 s, CPR duration was 168 ± 115 s.

**Table 1.** Baseline values of CA groups and sham-operated groups.

| Valuable | Sham/CA -3h |  |  | Sham/CA -24h |  |  | Sham/CA -72h |  |  |
| --- | --- | --- | --- | --- | --- | --- | --- | --- | --- |
|  | Sham | CA | *p*-value | Sham | CA | *p*-value | Sham | CA | *p*- |
| Gender(male) | 4 | 3 | >0.99 | 5 | 4 | >0.99 | 5 | 5 | >0.99 |
| Surviving rate(%) | 100(8/8) | 50(8/16) | 0.022 | 100(8/8) | 31(8/26) | 0.001 | 89(8/9) | 27(8/30) | <0.001 |
| Body weight(g) | 364±31 | 346±24 | 0.217 | 335±36 | 348±30 | 0.473 | 329±2 | 349±28 | 0.115 |
| Heart rate(beat/min) | 217±29 | 200±40 | 0.341 | 197±25 | 187±32 | 0.515 | 222±3 | 200±46 | 0.296 |
| MAP (mmHg) | 73±18 | 69±21 | 0.693 | 72±18 | 73±16 | 0.899 | 76±13 | 74±21 | 0.822 |
| RR(beat/min) | 73±17 | 83±19 | 0.306 | 77±13 | 70±12 | 0.347 | 90±18 | 77±14 | 0.131 |

MAP: Mean Arterial Pressure; RR: Respiratory Rate; CA: Cardiac Arrest

### Percentage of T cells and subsets

As shown in **Table 2**, compared with sham-operated group, the percentage of CD3+T cell in the CA group was decreased at 3 h post-ROSC and the differences were more significant at 24 and 72 h post-ROSC. No significance in the percentage of CD4+ and CD8+ lymphocyte subsets in the peripheral blood was observed between CA and sham-operated groups. Consistently, the ratio of CD4+/CD8+ have no significant difference in two groups.

**Table 2.** Comparison of T cells and subsets in the sham-operated groups and CA groups.

| Valuable | Sham/CA-3h |  |  | Sham/CA -24h |  |  | Sham/CA -72h |  |  |
| --- | --- | --- | --- | --- | --- | --- | --- | --- | --- |
|  | Sham | CA | *p*-value | Sham | CA | *p*-value | Sham | CA | *p*-value |
| CD3<sup>+</sup> ( % ) | 69.06±9.93 | 61.78±3.60 | 0.083 | 42.08±10.50 | 29.20±5.94 | 0.009 | 46.54±9. | 31.65±10.92 | 0.034 |
| CD4<sup>+</sup> ( % ) | 69.66±6.04 | 70.45±8.80 | 0.838 | 61.73±7.91 | 64.25±6.24 | 0.490 | 66.33±5. | 63.96±6.52 | 0.320 |
| CD8<sup>+</sup> ( % ) | 27.68±6.94 | 27.58±8.96 | 0.980 | 35.73±7.09 | 33.15±5.76 | 0.439 | 32.00±5. | 32.73±5.24 | 0.642 |
| CD4<sup>+</sup>/CD8<sup>+</sup> | 2.71±0.90 | 2.85±1.10 | 0.779 | 1.83±0.57 | 2.03±0.61 | 0.503 | 2.15±0.5 | 2.02±0.50 | 0.580 |

CA: Cardiac Arrest

### Plasma inflammatory cytokine levels

Compared with sham-operated group, the plasma concentrations of TNF-α in CA groups were noticeably higher at 3 and 24 h post-ROSC. However, at the three time points, there was no significant difference between sham-operated groups and CA groups (**Fig 1A**). In addition, the concentrations of IL-6 and IL-10 in CA groups were shown to achieve peak levels at 3 h post-ROSC, which was significantly higher than sham-operated at 3 h (128.74 (81.14-245.08) pg/ml vs 43.12(31.15-51.93) pg/ml, *p* = 0.001 and 252.80 (171.90-404.39) pg/ml vs 123.63 (73.39-212.52) pg/ml, *p* = 0.015) and at 24 h (36.56 (35.10-58.88) pg/ml vs 28.79(8.48-32.32) pg/ml, *p* = 0.005 and 69.50 (40.56-104.14) pg/ml vs 25.00(22.25-28.38) pg/ml, *p* < 0.001) (**Fig 1B, 1C**). In the following 2 days, the levels of IL-6 and IL-10 gradually decreased and returned to concentrations comparable with those found in sham-operated at 72 h (**Fig 1B, 1C**).

**Figure 1.**
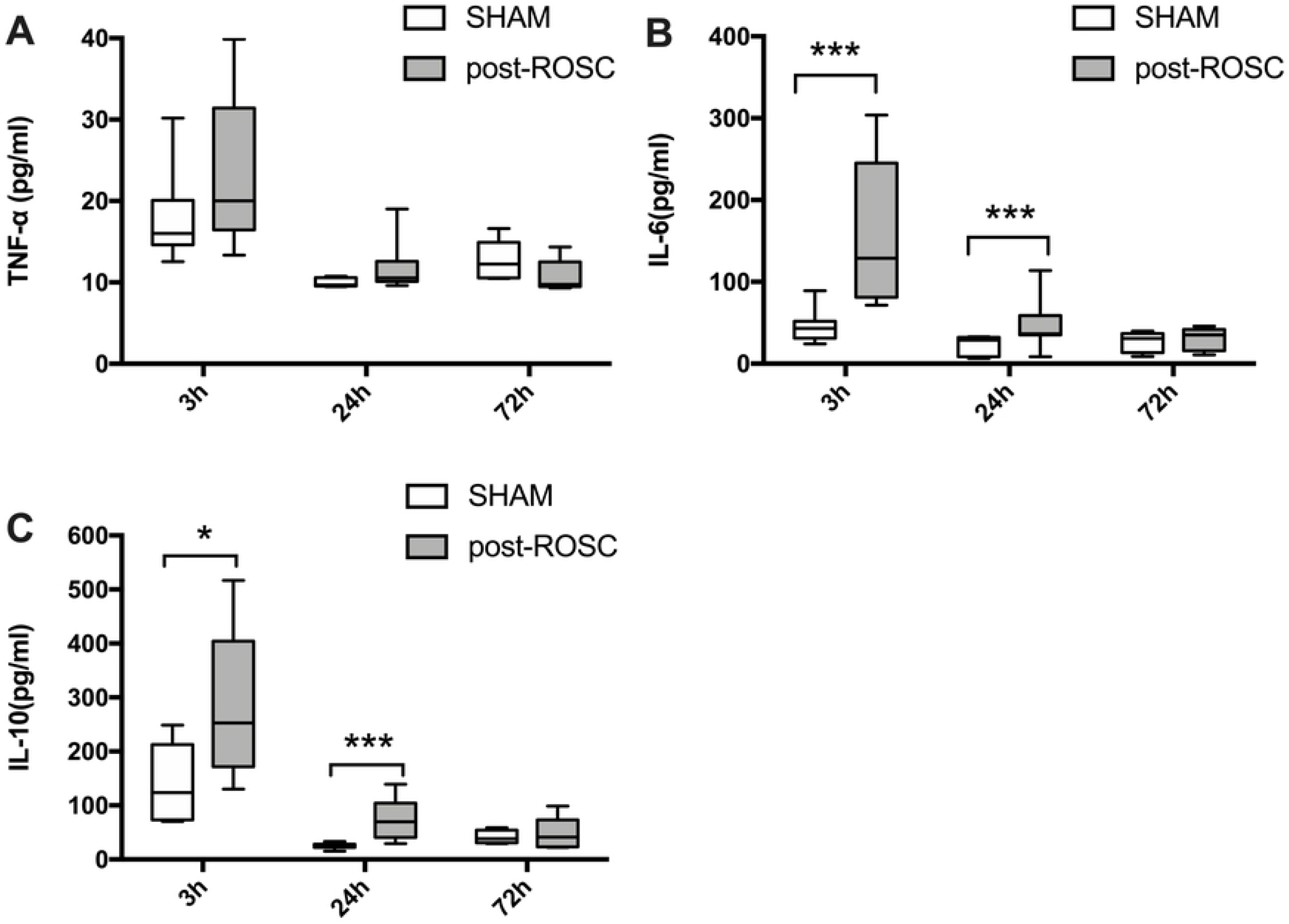
Levels (pg/mL) of the inflammatory cytokines tumor necrosis factor alpha (TNF)-α, interleukin (IL)-6 and IL-10 between CA groups and sham-operated groups at three time points. A, Levels of TNF-α; B, Levels of IL-6; C, Levels of IL-10. * p < 0.05, *** p < 0.001

### The co-stimulatory and co-inhibitory molecules expressed on CD4+ T cells

CD28, the co-stimulatory molecule, was demonstrated to be expressed on the surface of all CD4+ resting T cells. At the three time points, the expression of CD28 in the CA groups was obviously higher than that of in the sham-operated groups. Especially, the difference was particularly remarkable at 24 h post-ROSC (85.36 ± 22.46% vs 58.08 ± 24.95%, *p* = 0.037) (**Fig 2A**). Such changes also happened in the expression of CTLA-4 (3.66 (1.92-9.96) % vs 0.41 (0.08-0.79) %, *p* = 0.003)) and PD-1 (58.25 (44.6–90.68) % vs 3.05 (0.69-8.55) %, *p* < 0.001), the co-inhibitory molecules only expressed when T cells activated (**Fig 2B, 2C**).

**Figure 2.**
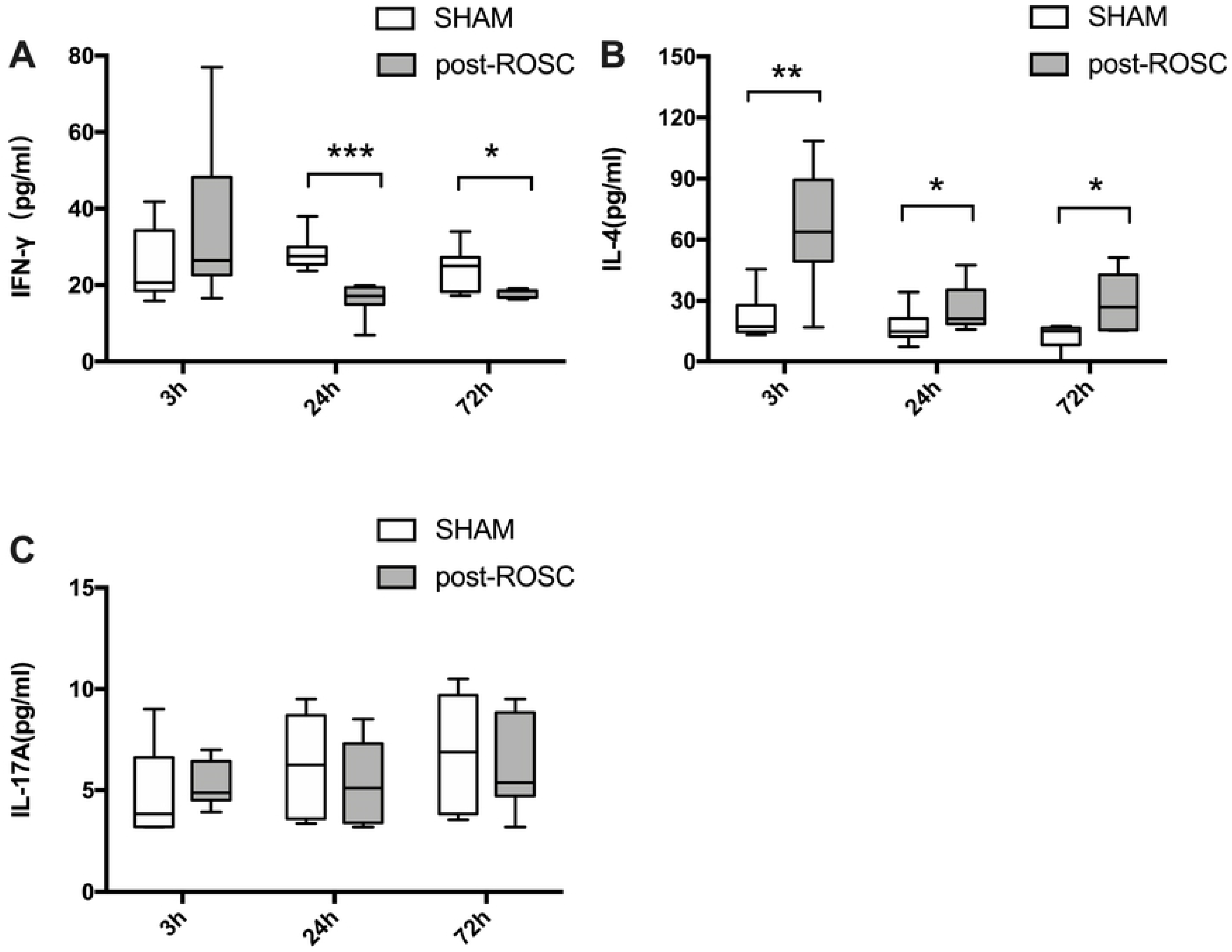
The percentage of CD28, CTLA-4, and PD-1 expressed on CD4+ T cells of CA groups and sham-operated groups at three time points. A, CD28; B, CTLA-4; C, PD-1 * p < 0.05, ** p < 0.01, *** p < 0.001.

### ELISA analysis of IFN-γ, IL-4, and IL-17A levels

IFN-γ, IL-4, and IL-17A are stated to be principally secreted by Th1, Th2 and Th17, respectively. Thus, these cytokine levels were measured to reflect the differentiate functions of T cells. In this study, it was found that IFN-γ was significantly decreased in CA groups compared with sham-operated groups at 24 h (16.25 ± 4.14 pg/ml vs 28.44 ± 4.37 pg/ml, *p* < 0.001) and 72 h (18.02 ± 1.00 pg/ml vs 24.13 ± 5.64 pg/ml, *p* = 0.018) (**Fig 3A**), while IL-4 was markedly increased (3 h: 63.99 (49.33-89.47) pg/ml vs 17.19 (14.67-27.79) pg/ml, *p* = 0.002; 24 h: 21.13 (18.52-35.16) pg/ml vs 14.80 (12.27-21.37) pg/ml, *p* = 0.028; 72 h: 26.89 (15.51-42.67) pg/ml vs 15.06 (8.22–16.70) pg/ml, *p* = 0.021) (**Fig 3B**). On the other hand, no significant difference was observed in IL-17A level at the three time points (**Fig 3C**).

**Figure 3.**
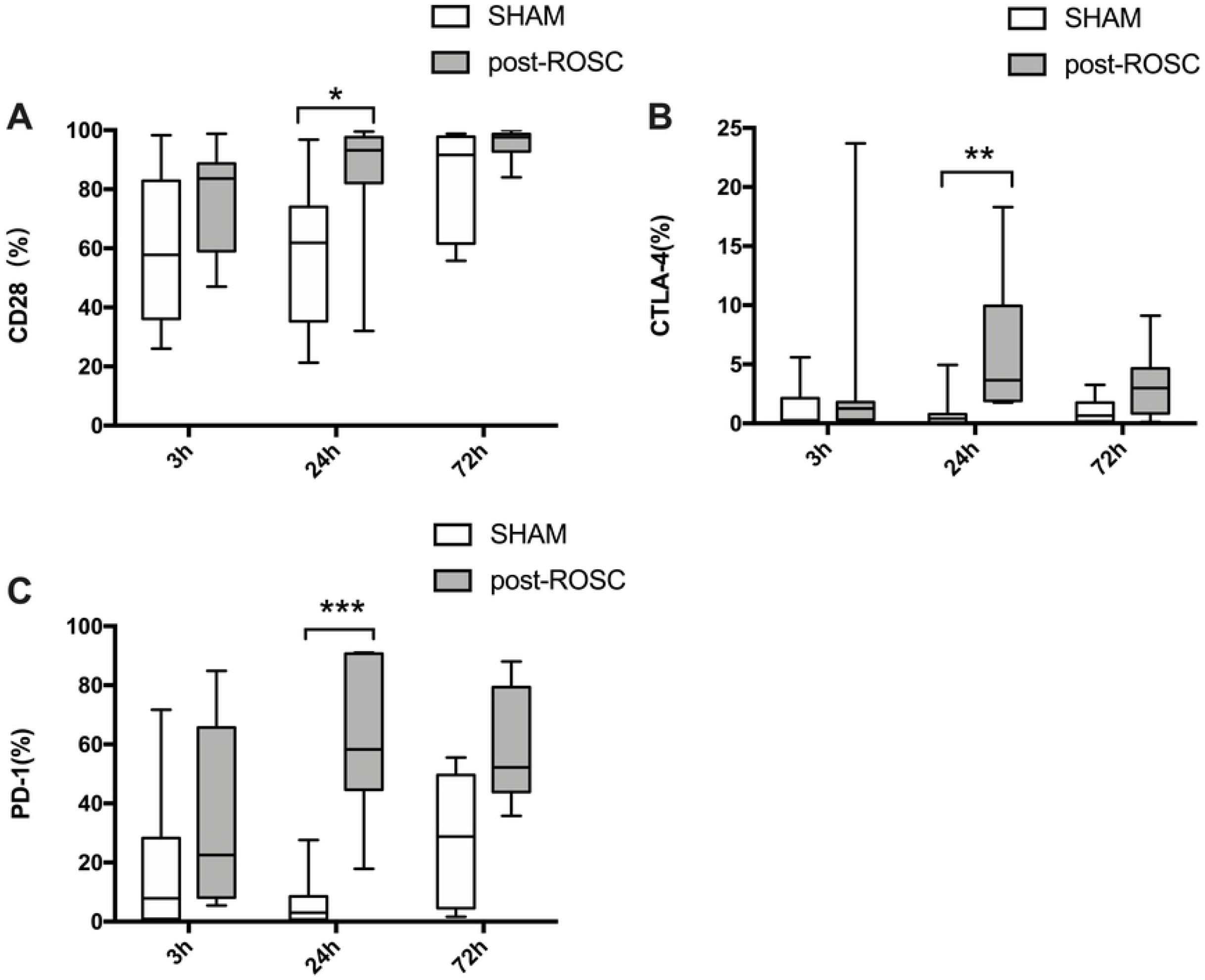
Levels (pg/mL) of the interferon γ (IFN-γ), interleukin (IL)-4 and IL-17A in plasma of CA groups and sham-operated groups at three time points. A, Levels of IFN-γ; B, Levels of IL-4; C, Levels of IL-17A. * p < 0.05, ** p < 0.01, *** p < 0.001.

## Discussion

To the best of our knowledge, the current study is the first time to evaluate the expression levels of CD28, CTLA-4 and PD-1 on the T cells of rats after cardiac arrest. The results suggest that CA rats show signs of immunoparalysis and the function of T cells is partially impaired in the post-ROSC rats. The impaired function manifested in the decreased percentage of T cells, the up regulated co-inhibitory molecules and the shift from Th1 to Th2 in blood.

During asphyxia, the hypoxemia and hypercapnia developed immediately, accompanied by transient tachycardia, elevated blood pressure and systemic vascular resistance, and the body has suffered the most severe shock with an interrupted transportation of oxygen and metabolites. After 5 min asphyxia, the body is demonstrated to present as bradycardia, hypotension, low cardiac output, pulmonary hypertension, tissue hypoxia and lactic acidosis, leading to the following CA [16]11. Although the hemodynamic, respiratory and systemic metabolic parameters could be restored after ROSC, hypoxia would persist in the tissues, especially the internal organs[16]. There is also an acute inflammatory response during cardiopulmonary resuscitation, not only the release of proinflammatory cytokines (such as IL-1, IL-6, TNF-α), but also the large number of inflammatory inhibitors (such as IL-1Ra, IL-4, IL-10 etc.), causing SIRS similar to sepsis [7, 17, 18]. These cytokines interact with each other and participate in the immune response and regulation of the organism. The observed high levels of inflammatory cytokines in the current study demonstrated a systemic inflammatory response syndrome (SIRS) was also developed during the first 24 h after CA.

CD3 is a characteristic marker on mature T cell surface, which could be mainly divided into CD4+T cell and CD8+T cell according to the expression of CD4 or CD8. Previous studies in trauma patients have confirmed that patients with reduced numbers of T cells have a higher risk to develop secondary infections [19]. In the current study, it was exhibiting that the decrease of T cells percentage was also observed after CA, while the percentages of CD4+ and CD8+ had no remarkable change. In addition, CD4+ lymphocytes were lower in the post-ROSC group at 12 and 24 h after ROSC, which was related to CA induced T cells death or apoptosis. The numbers of T cells, including CD4+ and CD8+, were therefore reduced and the proportions remained normal [9, 10].

The activation of T cells requires dual signal stimulation, during which the first signal is provided by the TCR/CD3 antigen peptide. T cells and adhesion molecules on antigen-presenting cell surface subsequently provide the second signal, which is called co-stimulatory molecules. CD28 is indicated to be the most important co-stimulatory molecules expressed on T cells [20]. The level of CD28 expression on the T cells in severely infected patients is significantly lower, which is an independent risk factor for increased mortality, and also a good marker to predict the outcome of septic shock patients [21]. In our study, all animals had a good prognosis (i.e. survive from cardiac arrest), which may explain the high expression of CD28.

The Cytotoxic T lymphocyte-associated antigen (CTLA)-4 is a co-inhibitory molecule expressed mainly on the surface of activated T cells and is highly homologous to CD28 [20]. The main function of CTLA-4 is to inhibit he binding of CD28 to B7 and to limit the activation and proliferation of T cells [22]. Furthermore, CTLA-4/Ig fusion protein competitively inhibits CD28 and B7 binding, which can play a strong immune effect and has been used in clinical treatment [23]. In the animal model of sepsis, elevated expression of CTLA-4 was found and blocking CTLA-4 pathway could improve survival rate [24]. In our study, CTLA-4 in CA groups significantly increased at 3h, 24h and 72h. Especially at the 24h, the level of this co-stimulatory molecules reached its high peak. All these changes confirmed a similar change with that progression of sepsis studies, indicating T cells were in a state of inhibition after CA.

As a co-inhibitory molecule on activated lymphocytes, PD-1 attenuates T-cell responses and is important in the regulation of T-cell tolerance [21]. Previous studies have suggested that PD-1 plays an immune suppression role on patients with sepsis and on the mouse model of infection [23, 25]. The upregulated expression of PD-1 has been detected in septic patients and correlated with worse outcome [25]. Similarly, in the mouse sepsis model, the expression rate of PD-1 on the T cells was also increased. while blocked the signal pathway of PD-1 could increase the clearance rate of pathogenic microorganisms and rise the survival rate of the septic rats [23]. In the current study, PD-1 was constantly in a high expression state at different time point, suggesting that T cells may be in a state of sustained inhibitory. Further studies are required to investigate whether such high expression status could cause severe depletion of T cells as the findings in sepsis patients and its impact on prognosis [26].

Activated CD4+T cells briefly differentiate into three kinds of helper cells with different functions, namely, Th1, Th2 and Th17 cells [27]. The specific cytokines secreted by helper T cells are detected to reflect the different function. IL4 and IFN-γ concentrations were witnessed a significant difference in two groups in our study. A drift of Th1 to Th2 has been found in severe septic patients, which is associated with immune dysfunction [28]. Moreover, previous studies found a shift from Th2 to Th1 response in the myocardium in a porcine model of cardiac arrest, which is associated with the redistribution of circulating T lymphocyte subsets [10, 21]. While previous studies examined the immune response in the myocardium, the cytokine concentrations in the serum were also found a shift from Th1 state to Th2, which was similar to the findings in sepsis [28]. Additional studies are required to gain more insight in the mechanisms of redistribution of immune response. As an essential factor associated with innate immunity and adaptive immunity, inflammatory cytokine IL-17A plays an important role in innate immunity [29]. In the present study, IL-17A was slightly but not significantly elevated at three time points after CA, which is consistent with the results of Callaway et al [30], indicating that the differentiation process of CD4+T cells into Th17 may be unaffected after CA.

The current study also has several limitations. Firstly, arterial blood gas analysis was not performed and the dynamic change of vital signs was not recorded. Aside from the function change in the preliminary study, the mechanism of immunity alteration after CA are required to further investigate.

## Conclusions

In summary, the results of this study showed that the occurrence of cardiac arrest affected T cells function as illustrated by decreased percentage of T cells, the up regulated co-inhibitory molecules and the shift from Th1 to Th2 in blood. Furthermore, the expression levels of CD28, CTLA-4 and PD-1 on the T cells of rats were evaluated after cardiac arrest. These preliminary results should be confirmed in a large cohort of OHCA patients and the relationship between the function change of T cells and the prognosis requires to be further explored after CA.

## Acknowledgment

This study was supported by Shanghai Soong Ching Ling Foundation. The organization had no role in study, design, collection, analysis, and interpretation of data, in the writing of the report, or in the decision to submit the article for publication.

## Author Contributions

Conceptualization: Weiming Chen, Guoping Lu, Chunlin Xing

Methodology: Chunlin Xing, Xuemei Zhu

Project administration: Yang Chen, Chunlin Xing

Supervision: Guoping Lu, Weiming Chen

Writing – original draft: Chunlin Xing

Writing – review & editing: Weiming Chen, Guoping Lu

